# rasterdiv - an Information Theory tailored R package for measuring ecosystem heterogeneity from space: to the origin and back

**DOI:** 10.1101/2021.02.09.430391

**Authors:** Duccio Rocchini, Elisa Thouverai, Matteo Marcantonio, Martina Iannacito, Daniele Da Re, Michele Torresani, Giovanni Bacaro, Manuele Bazzichetto, Alessandra Bernardi, Giles M. Foody, Reinhard Furrer, David Kleijn, Stefano Larsen, Jonathan Lenoir, Marco Malavasi, Elisa Marchetto, Filippo Messori, Alessandro Montaghi, Vítězslav Moudrý, Babak Naimi, Carlo Ricotta, Micol Rossini, Francesco Santi, Maria J. Santos, Michael Schaepman, Fabian Schneider, Leila Schuh, Sonia Silvestri, Petra Šímová, Andrew K. Skidmore, Clara Tattoni, Enrico Tordoni, Saverio Vicario, Piero Zannini, Martin Wegmann

## Abstract

1. Ecosystem heterogeneity has been widely recognized as a key ecological feature, influencing several ecological functions, since it is strictly related to several ecological functions like diversity patterns and change, metapopulation dynamics, population connectivity, or gene flow.
2. In this paper, we present a new `R` package - `rasterdiv` - to calculate heterogeneity indices based on remotely sensed data. We also provide an ecological application at the landscape scale and demonstrate its power in revealing potentially hidden heterogeneity patterns.
3. The `rasterdiv` package allows calculating multiple indices, robustly rooted in Information Theory, and based on reproducible open source algorithms.

---

> ‘*I’ll be back* ‘ (A. Schwarzenegger, The Terminator, 1984)

## 1 Introduction

Ecosystem heterogeneity drives a number of ecological processes and functions such as species diversity patterns and change (Rocchini et al., 2018), metapopulation dynamics (Fahrig, 2007), population connectivity (Malanson and Cramer, 1999) or gene flow (Lozier et al., 2013). Heterogeneity has been defined in various ways in the scientific literature: (i) as the variation in space and time of qualitative and quantitative descriptors of an environ-mental variable of interest (Li and Reynolds, 1995); (ii) as the horizontal component of habitat variation (August, 1983; Grelle, 2003); (iii) as the spatially structured variability of the habitat (Ettema et al., 2002); or (iv) as within-habitat variability (Heaney, 2001; Hortal et al., 2009). In this paper, it will be considered as an umbrella concept representing the degree of non-uniformity in land cover, vegetation and physical factors (topography, soil, topoclimate and microclimate, Stein et al., 2014).

Landscape heterogeneity across different spatial extents and over different temporal periods can be inferred by applying algorithms based on remote sensing and spatial ecology (Skidmore et al., 2011; Schimel and Scheiner, 2019). Remotely sensed spectral heterogeneity measures of a landscape represent a valid alternative to categorical land cover maps, which, especially in the case of non-homogeneous and complex landscape configurations (e.g. mosaic of crops and semi-natural forests), might suffer from oversimplification when investigated through land cover classes (Rocchini et al., 2019; Da Re et al., 2019). Heterogeneous landscapes should present higher spectral heterogeneity values compared to more homogeneous landscapes within the same spatial extent (Rocchini and Ricotta, 2007). It follows that remotely sensed spectral heterogeneity can be profitably used to measure landscape heterogeneity in space and time in order to convey information on ecosystem processes and functioning (Schneider et al., 2017).

From this point of view, the development of Free and Open Source algorithms to measure and monitor (i.e. repeated measures over time) landscape or ecosystem heterogeneity from space would allow robust, reproducible and standardized estimates of ecosystem functioning and services (Rocchini and Neteler, 2012). Furthermore, their intrinsic transparency, community-vetoing options, sharing and rapid availability are also valuable additions and reasons to move towards open source options. Among the different open source software options, the `R` software is certainly one of the most used statistical and computational environment worldwide and different packages have already been devoted to remote sensing data processing for: (i) raster data management (`raster` package, Hijmans and van Etten (2020)); (ii) remote sensing data analysis (`RStoolbox` package, Leutner et al. (2019)); (iii) spectral species diversity (`biodivMapR` package, Féret and Boissieu (2020)); (iv) Sparse Generalized Dissimilarity Modelling based on remote sensing data (`sgdm` package, Leitao et al. (2017)); (v) entropy-based local spatial association (`ELSA` package, Naimi et al. (2019)); or (vi) landscape metrics calculation (`landscapemetrics` package, Hesselbarth et al. (2019)), to name just a few. Readers can also refer to https://cran.r-project.org/web/views/Spatial.html for the CRAN Task View on analysis of spatial data.

Nonetheless, currently no R package provides a flow of functions grounded in Information Theory and generalized entropy, incorporating abundance information for each informative value but also the relative numerical distance among the said values. In this paper we introduce the new `rasterdiv` R package, now available under the Comprehensive R Archive Network (CRAN, https://CRAN.R-project.org/package=rasterdiv), which provides such a set of functions’ throughput workflow. The aim of this manuscript is to briefly introduce the theory under the `rasterdiv` package and to provide an ecological example demonstrating its ability to measure several aspects of landscape or ecosystem heterogeneity.

## 2 Brief description of the theoretical frame-work on information theory based metrics

### 2.1 Shannon entropy

Shannon’s theory (Shannon, 1948), profoundly rooted in Eduard Boltzmann (Boltzmann, 1872) studies, is a solid basis for calculating landscape heterogeneity and it has been widely used in several ecological applications (refer to Vranken et al., 2014 for a review).

Given a sample area with *N* pixel values and *p*_*i*_ relative abundances for every *i* ∈ {1, …, *N}*, in decreasing order, the Shannon index is calculated as:

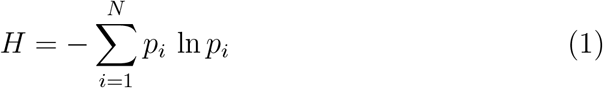

When applying such indices to remotely sensed data, the image is divided into small chunks of the whole image, commonly defined as ‘kernels’, ‘windows’ or ‘moving windows’ (Box 1). These terms will be used throughout this manuscript to relay the local scale of analysis.

#### Box 1 Description of the moving window approach

Given a raster layer, ecologists usually split it into small chunks, called moving windows. To estimate the value of the previously presented indices, we first select *l* an odd number, which will correspond to the length of the squared window. With this choice, we have a central entry in the window, i.e. the 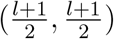 entry, which will be placed as a mask, in position (1, 1) over our raster. The index is therefore computed only with the values which the window covers. Notice that with this choice we will have some missing values, which will not contribute in the index computation. The obtained index value is stored in position (1, 1) in the output raster. The following step is moving the window so that its central entry is over the entry (1, 2) of the raster. The index value computed is stored in the corresponding position of the output raster, i.e. in (1, 2). We proceed in this way until the last entry of output raster is filled. This technique, visually presented in Figure 4, is extremely popular in ecology since it is quite simple, robust and powerful to simulate a biological boundary. Note that this technique of computing index has a suitable algorithmic structure.

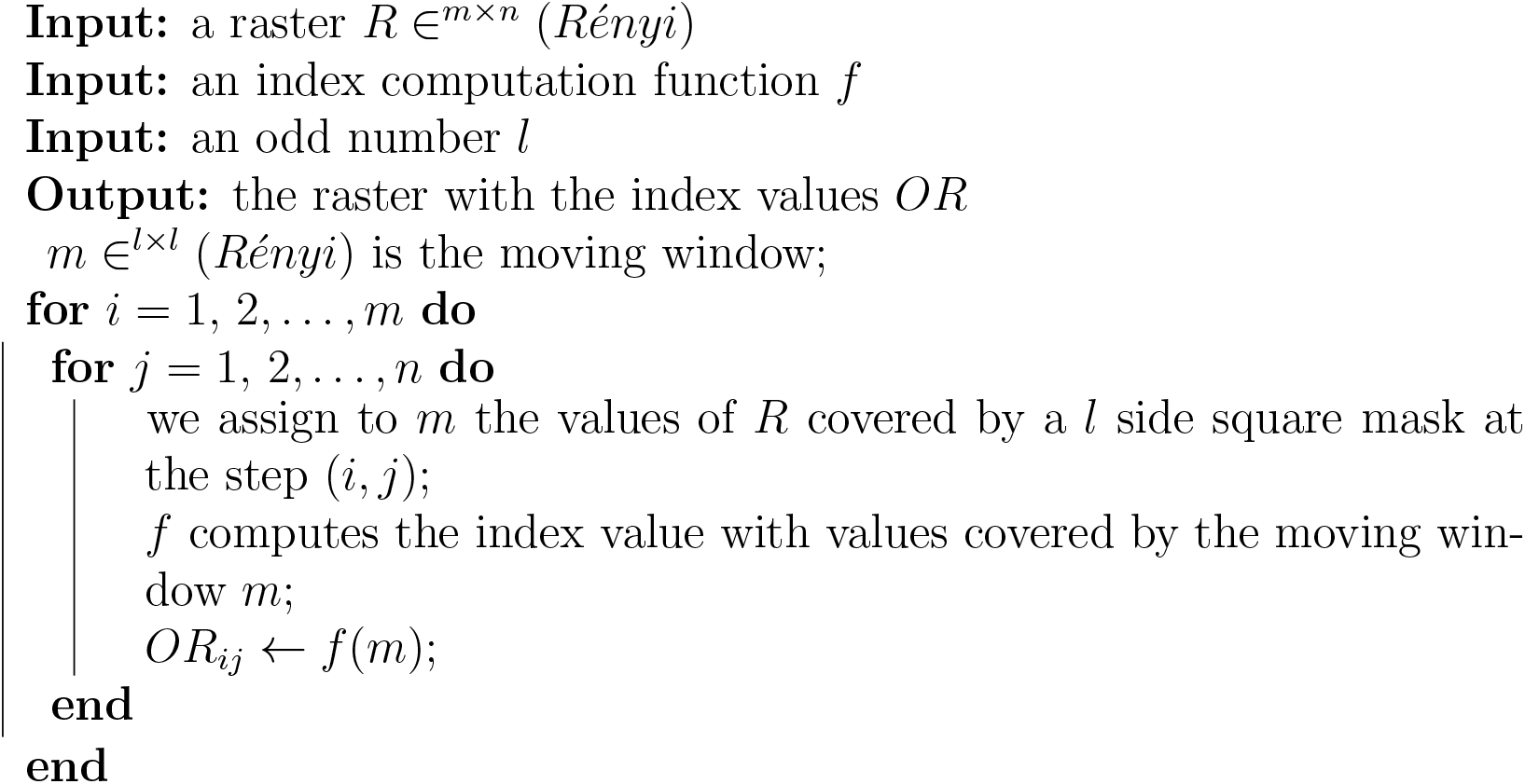

In the Shannon index the relative abundance of pixel values (e.g. reflectance values, vegetation indices) is considered. The higher the richness and turnover, the higher will be the equitability of values and thus the Shan-non index.

### 2.2 Rényi generalized entropy

Any point descriptor of information heterogeneity, like the previously cited Shannon’s *H* is not able to describe the whole potential spectrum of heterogeneity as it usually measures only one part or component of heterogeneity (e.g. richness, evenness, nestedness, etc.). Hence, no single measure can be used to represent such a wide spectrum (Gorelick, 2011a; Nakamura et al., 2020).

Rényi (1970) proposed a method to generalize entropy measurements in just one formula, changing one parameter, called *alpha* in its original formulation.

Given a sample area with *N* pixel values and *p*_*i*_ relative abundances for every *i* ∈ {1, …, *N*}, the Rényi entropy index is:

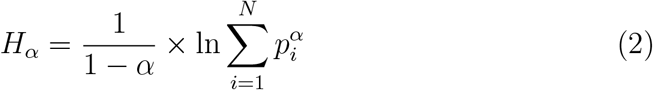

Changing the parameter *α* will lead to different indices starting from the same formula (Hill, 1973). As an example, when *α*=0, *H*_0_ = ln(*N*) where *N* =richness, namely the maximum possible Shannon index (*H*_*max*_). In practice, with *α* = 0, all the spectral values equally contribute to the index, without making use of their relative abundance. For *α* → 1, the Rényi entropy index will equal Shannon *H*, according to the l’Hôpital’s rule (a mathematical proof is provided in Appendix 1), while for *α*=2 the Rényi entropy index will equal the ln(1/*D*) where *D* is the Simpson’s dominance (Simpson, 1949). The theoretical curve relating the Rényi entropy index and *α* is a negative exponential, i.e. it decays until flattening for higher values of *α*, where the weight of the most abundant spectral values is higher with small differences among the attained heterogeneity maps (Ricotta et al. (2003a)). Besides the Rényi’s generalized entropy, the `rasterdiv` package includes generalized metrics as the Hill’s numbers (Hill, 1973).

### 2.3 Rao’s *Q* heterogeneity index

The previously described metrics have no dimension. In other words, they do not consider the relative difference among pixel values but just the presence of a different class. For example, having *A* = (1, 2, 3, 4, 5, 6, 7, 8, 9) and *B* = (1, 10^2^, 10^3^, 10^4^, 10^5^, 10^6^, 10^7^, 10^8^, 10^9^) as two theoretical arrays containing values that are different from each other, the Shannon index will always be maximum, i.e. *H* = log(9) = 2.197225, despite the relative numerical distance between pairs of values.

In remote sensing this is a critical point since contiguous zones of a satellite image might have similar (but not strictly equal) reflectance values. For instance, the variability of a homogeneous surface, e.g., a woodland patch or a water area, would be overestimated if spectral distances among values are not considered in the calculation.

The Rao’s Quadratic heterogeneity measure (hereafter Rao’s *Q*, Rao (1982)) can be applied to overcome this issue, considering both relative abundances and spectral distances among pixel values in the calculation.

Given the values of different pixels *i* and *j*, the Rao’s *Q* considers their pairwaise distance *d*_*ij*_ as:

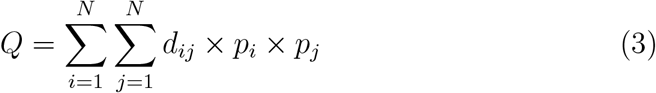

Accordingly, an array with different but spectrally near values will convey a high Shannon’s *H* but a low Rao’s *Q*. Conversely, an array with different and distant spectral values will convey both a high Shannon’s *H* and a high Rao’s *Q*.

Given a 3×3 pixels matrix M: 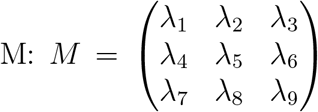where *λ* is the reflectance value for every pixel in a single 8-bit band (256 possible values), a pairwise distance matrix *M*_*d*_ is derived for all pixel values:

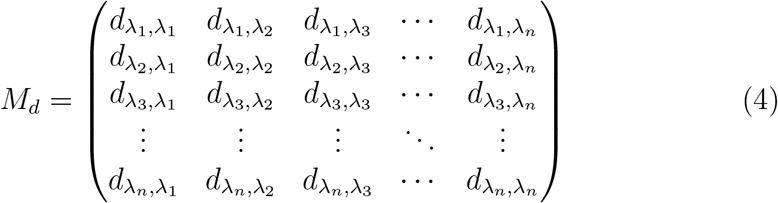

Then, according to Equation 3, Rao’s *Q* is obtained as the sum of every pairwise distance multiplied by the relative abundances of all the pairs of pixels in the analyzed image *d* ×(1*/N* ^2^). Strictly speaking, Rao’s *Q* can be defined as the expected difference in reflectance values between two pixels drawn randomly with replacement from the evaluated set of pixels.

It is possible to construct the distance matrix for several dimensions (layers) in order to consider multiple bands at a time and calculate Rao’s *Q* in a multidimensional (multi-layers) system.

To illustrate the benefit of the functions provided in the `rasterdiv` package, we propose to apply it on an ecological study case in the following section.

## 3 Application: the ecosystem heterogeneity of the Ötzi area

In order to show the capabilities of the `rasterdiv` package, we decided to focus on one of the geologically and biologically most diverse mountain regions worldwide: the Similaun and Ortles glaciers in Italy. This region is not only important for its rich geobiodiversity but also for its fascinating archeological history, also due to an incredible anthropological discovery of the early nineties: the famous Ötzi Tyrolean iceman (Keller et al., 2012) The study area we are focusing on for applying the `rasterdiv` package is included between the Ortles and the Similaun glaciers, in the Alps of northern Italy (Figure 1). Below, we provide a step by step tutorial which can be reproduced for any area and by every researcher worldwide. The only required input data are satellite images.

**Figure 1:**
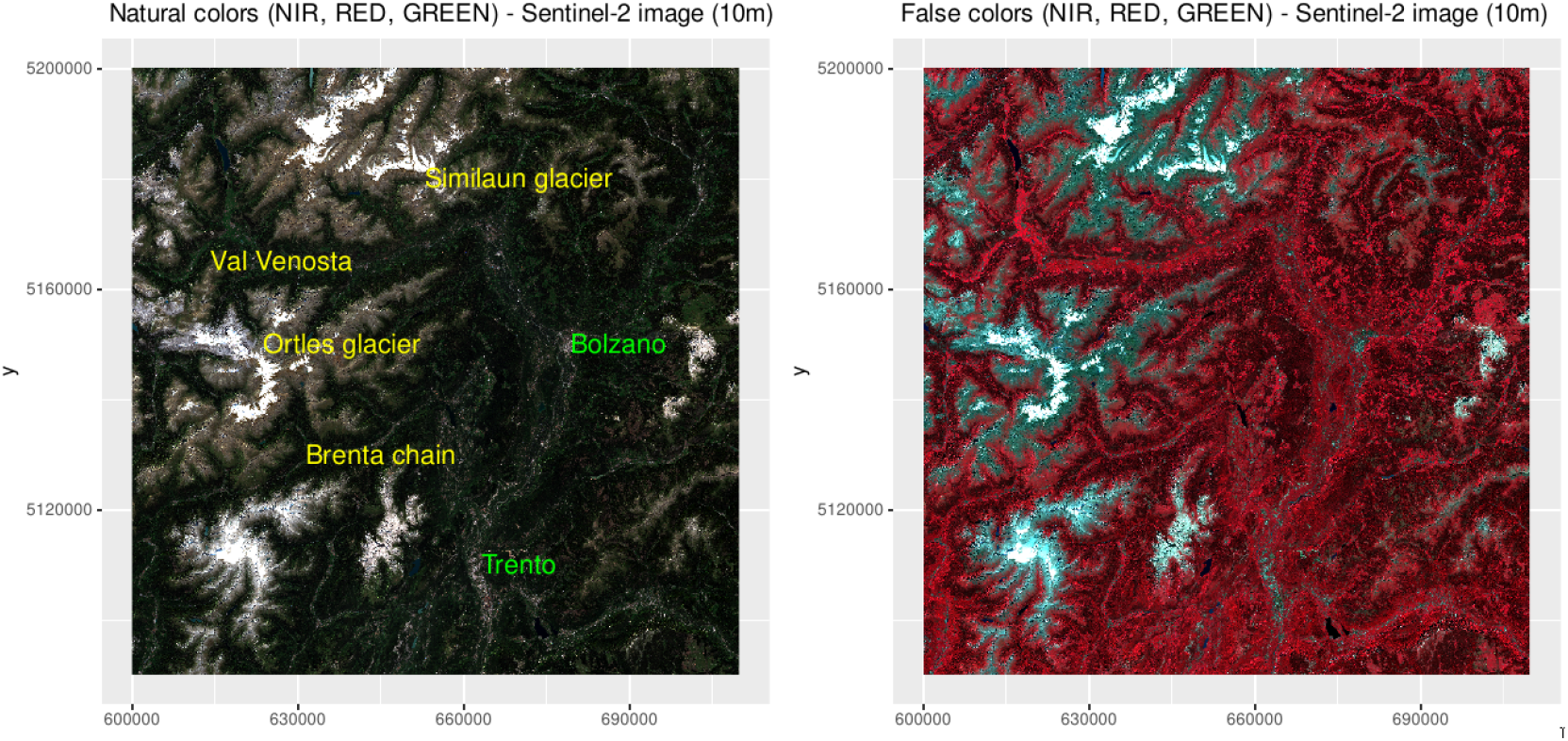
The Ötzi area in the northern italian Alps, used for building an ecological example of the application of the `rasterdiv` package starting from a Copernicus Sentinel-2 image, represented by an RGB space in natural (red, green, blue) and false (near infrared, red, green) colors. Coordinates are in the UTM (WGS84, zone 32N) reference system. High resolution figure available upon request.

We used Copernicus Sentinel-2 data at a spatial resolution of 10 m (Figure 1) acquired on May 9th 2020. Once the satellite image was downloaded from the Copernicus Open Access Hub **(https://scihub.copernicus.eu/)**, we computed NDVI and rescaled it at an 8-bit radiometric resolution. Hence, NDVI values were used as the input information (values) to compute the ecosystem heterogeneity indices described in the former section. More specifically, based on the NDVI raster grid used as input object, we ran a set of functions provided in the `rasterdiv` package and written in Box 2.

### Box 2 Code used under rasterdiv

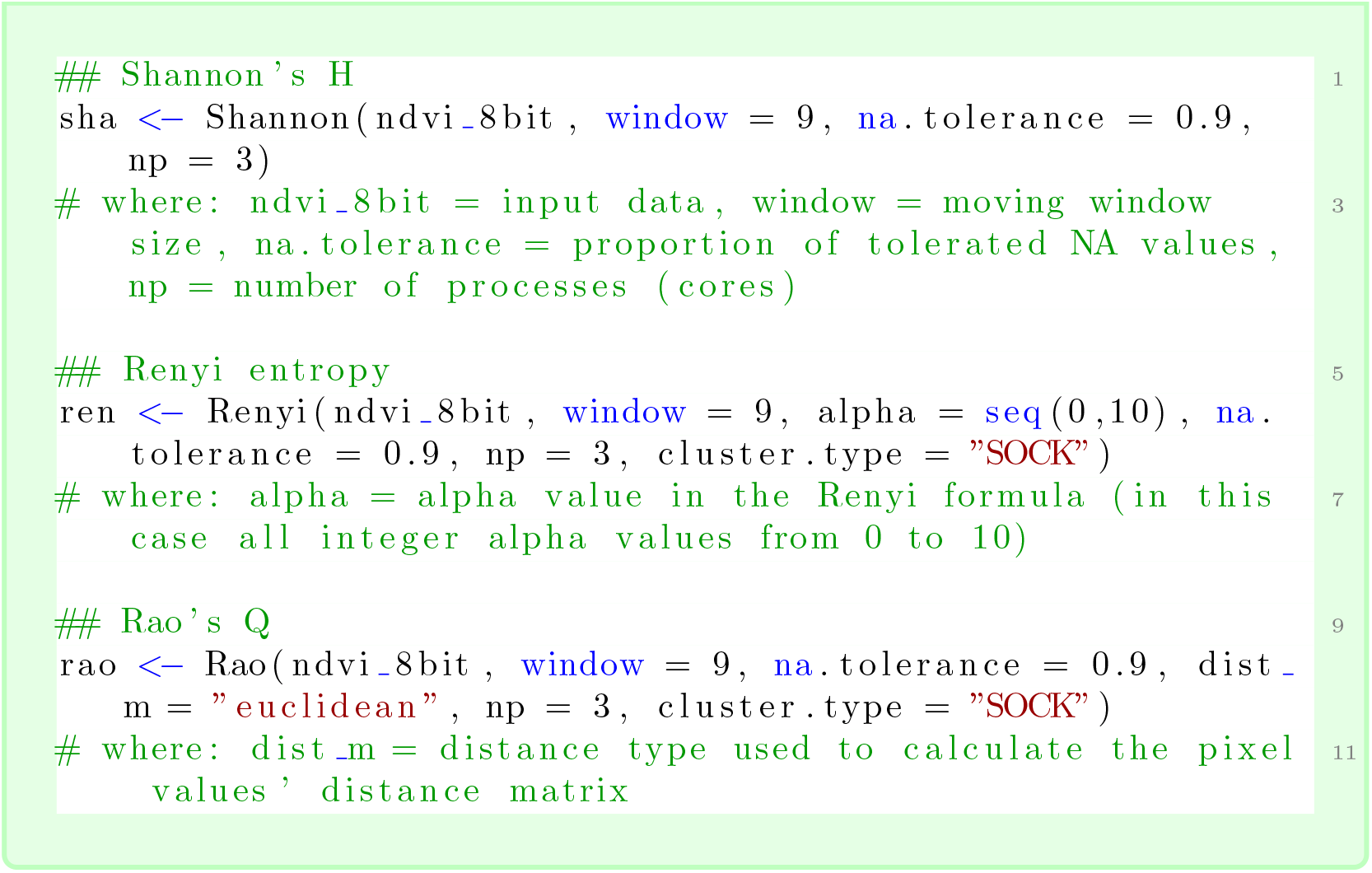

We applied the code written in Box 2, using a moving window of 9×9 pixels, to both: (i) the Similaun glacier upper area, mainly covered by alpine conifer woodlands (dominated by *Picea abies, Larix decidua, Abies Alba*) and rocks, as well as (ii) to the valley bottom (Val Venosta), a human-dominated landscape devoted to agricultural areas and small urban villages.

Concerning the Similaun glacier area, the Shannon index showed medium to high values everywhere, including areas with almost homogeneous rock cover (i.e. alpine habitat) as well as areas with homogeneous tree cover (i.e. coniferous forest habitat) (Figure 2). This is due to the fact that Shannon’s *H* does not take into account the distance among pixel values but only the relative abundance of each value within the moving window of 9×9 pixels. In this case, NDVI values showed subtle differences among each others, especially in homogeneous areas and even when rescaled at 8 bit (namely 256 possible integer values).

**Figure 2:**
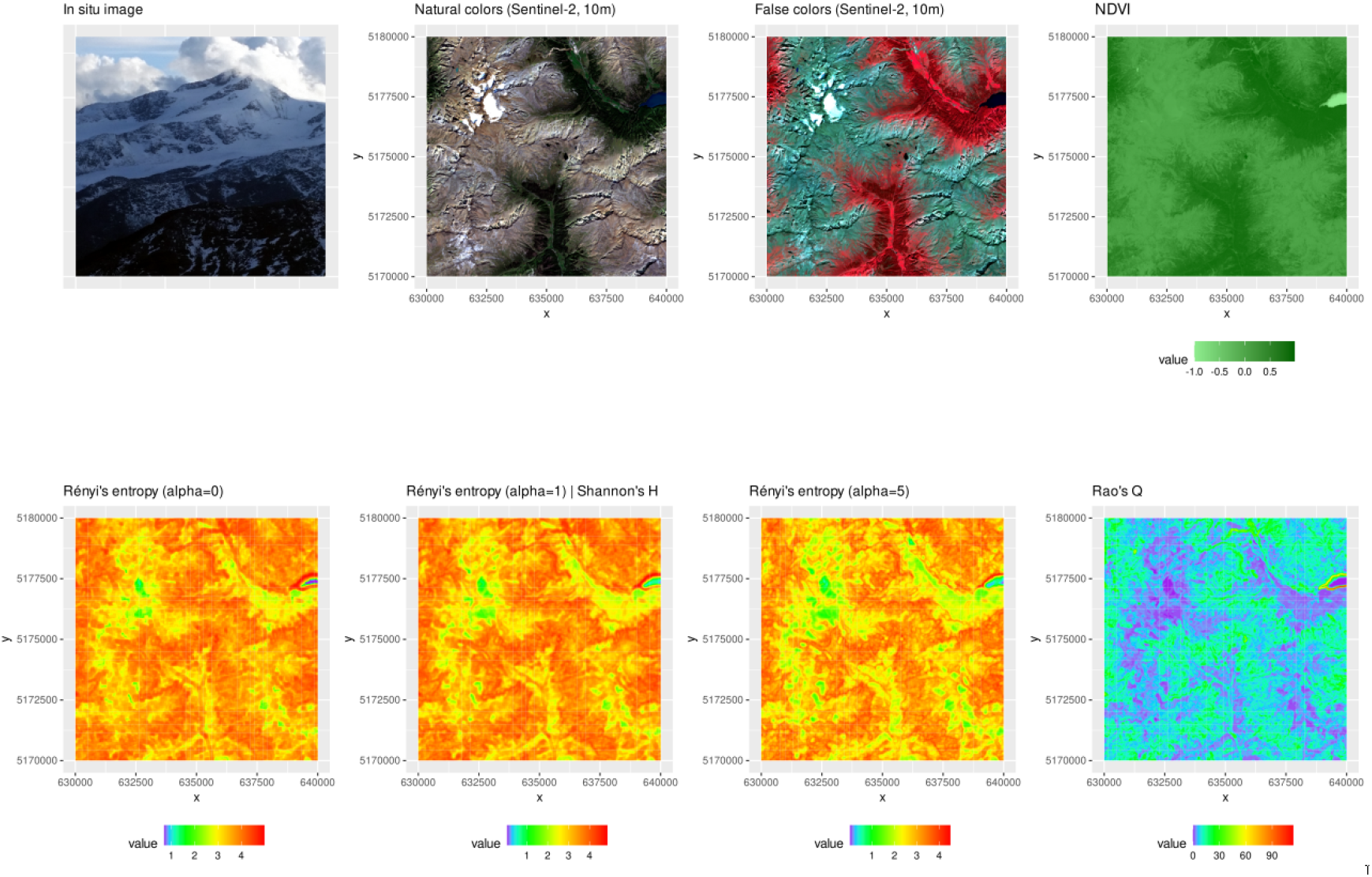
The first row of the figure represents the area under study, namely the Similaun glacier as a subset of the study area in Figure 1. NDVI is shown in the natural range but it was rescaled to 8-bit before heterogeneity computation. Then, different metrics were applied: the Rényi’s entropy index, the Shannon’s *H* (corresponding to the Rényi’s entropy with *α* = 1), and the Rao’s *Q*. Coordinates are in the UTM (WGS84, zone 32N) reference system. High resolution figure available upon request.

Low heterogeneity values were found in areas with snow cover; in that case, the values of the neighboring pixels are so similar that they lead to a low Shannon’s *H* for the focal pixel centered on the moving window. The saturation of high values of heterogeneity was apparent when considering Rényi’s entropy at low alpha values (Equation 2). This is related to the aforementioned (see section ‘2.2 Rényi generalized entropy’) negative exponential curve relating the value of this index with respect to alpha, shown in Ricotta et al. (2003a). The result is a map with saturated values of hetero-geneity. This effect is softening when increasing the alpha parameter. The `rasterdiv` package allows accounting for several indices at a time to avoid heterogeneity saturation effects, for instance by considering distance among pixel values besides relative abundance of each value. More specifically, running the Rao’s *Q* function coded in `rasterdiv` allows to circumvent this issue of saturated values of heterogeneity (refer to the bottom line of Figure 2). In the Similaun glacier area, the homogeneous cover of spruce forests is better reflected by the Rao’s *Q* index as it better contrasts against the geological heterogeneity of the upper alpine belt. Maximum Rao’s *Q* values were found at the interface between water (i.e. alpine lakes) and the surrounding vegetation and rocks, representing an interesting ecotone area (see the upper right corner of the last panel in Figure 2).

The `rasterdiv` package, thanks to a combination of functions rooted in Information Theory, thus helps to reveal hidden spatial patterns of heterogeneity, and allows measuring ecosystem heterogeneity related to both biotic and abiotic components. By doing so, the `rasterdiv` package better discriminates among ecosystem features and functions constituting the landscape.

In the valley area (Val Venosta), this phenomenon was even more pronounced (Figure 3) due to the higher spatial heterogeneity of the landscape matrix under study with patches of small agricultural fields mixed with water bodies (e.g., the Adige river in the middle of the valley), resulting in very high values for both Rényi (alpha=0) and Shannon indices. Low to medium values on the north- and south-facing slopes, indicate grasslands and broadleaf forests. Again, the Rao’s *Q* index helped to better discriminate among these areas and among different land uses. Indeed, by relying on the relative numerical distance among the 8bit-NDVI pixel values, it can differentiate between areas with low to medium heterogeneity values (blue and light blue colors in Figure 3), i.e. grasslands and broadleaf forests, and areas with medium to high heterogeneity values, i.e. upper mountain rocks at the interface with the treeline and with alpine lakes, as well as riparian vegetation besides the Adige river in the lower part of the valley.

**Figure 3:**
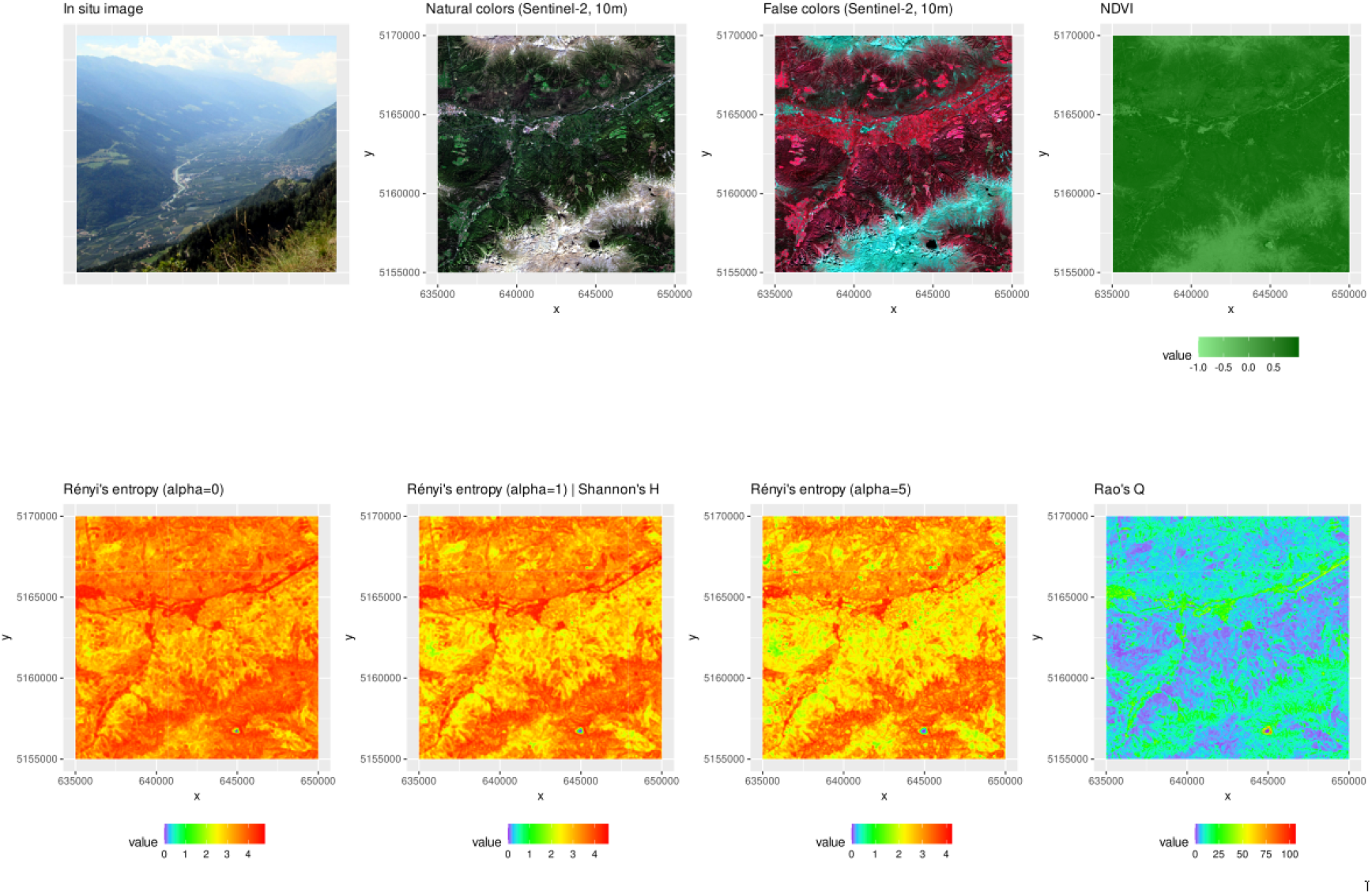
The first row of the figure represents the area under study, namely the Val Venosta, a subset at a lower elevation with respect to the Similaun glacier of Figure 2, and thus with higher human impact. NDVI is shown in the natural range but it was rescaled to 8-bit before heterogeneity computation. Then, different metrics were applied: the Rényi’s entropy, the Shannon’s *H* (corresponding to the Rényi’s entropy with *α* = 1), and the Rao’s *Q*. Coordinates are in the UTM (WGS84, zone 32N) reference system. High resolution figure available upon request.

**Figure 4:**
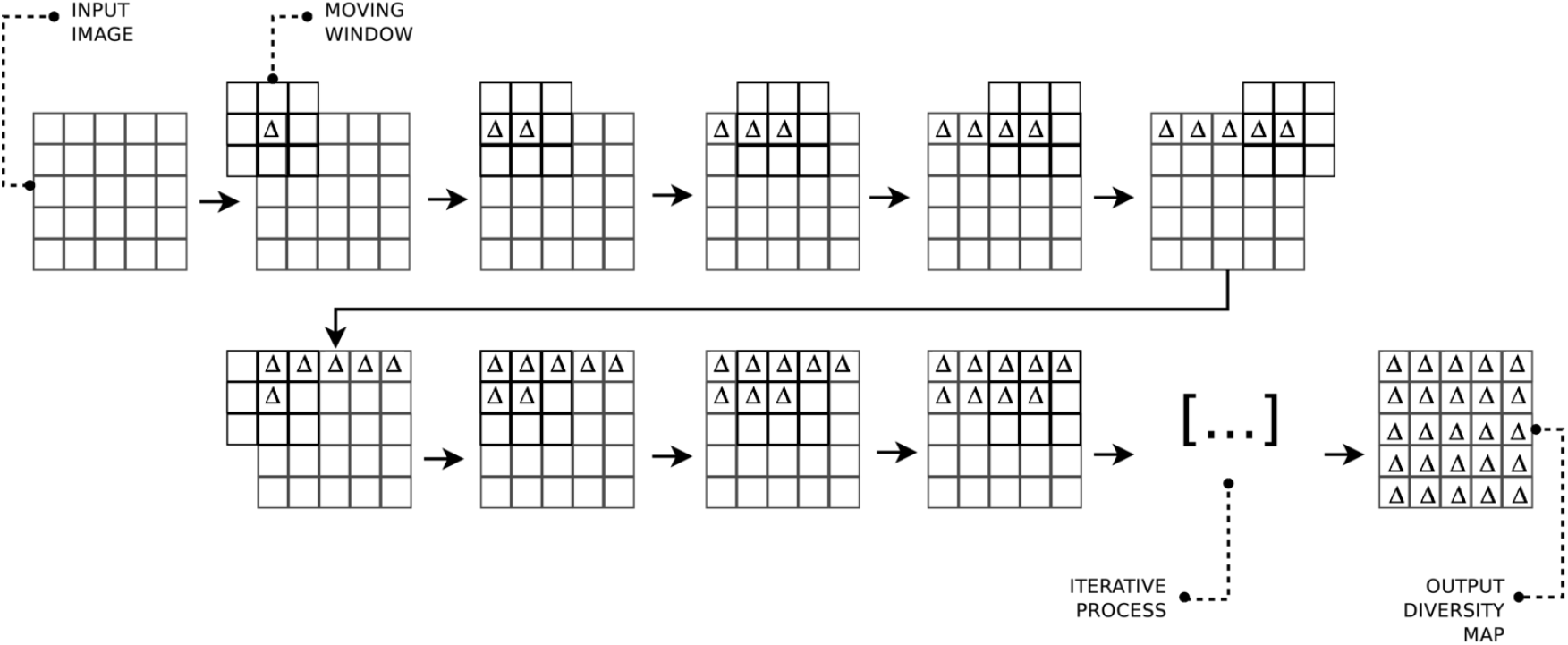
The moving window technique for the computation of diversity indices in the `rasterdiv` package, redrawn from Rocchini et al. (2013).

With this study case focusing on a very heterogeneous area in the Alps, we illustrated two main critical components of the `rasterdiv` package: (i) how it allows to measure multiple indices simultaneously, and (ii) how it can help detecting otherwise hidden patterns of heterogeneity in the landscape.

For the sake of clarity and to avoid a catalogue-like article, we did not present all the indices that can be calculated by the `rasterdiv` package. Instead, we decided to showcase a few, to illustrate to ecologists the high potential of the `rasterdiv` package for applications in landscape ecology, macroecology, biogeography and the analysis of spatiotemporal dynamics in general. We refer to the manual of the package **(https://cran.r-project.org/web/packages/rasterdiv/index.html)** and its vignettes to find additional metrics and examples related to the package.

## 4 Discussion

In this paper, we provide an ecological overview of new R package `rasterdiv`. The overarching rationale for proposing this new R package is to present a set of methods and ready-to-use functions for calculating landscape heterogeneity metrics from space (e.g. satellite images), as well as from airborne or ground-based devices, to monitor and analyze, among other things, bio-diversity change, habitat fragmentation and land use and cover changes.

The importance of computing continuous spectral heterogeneity measures from satellite-borne or airborne sensors to better discriminate among the various components in the landscape has been highlighted in several studies (Karlson et al., 2015; Godinho et al., 2018; Ribeiro et al., 2019; Doxa et al., 2020). Nevertheless, caution is recommended when making use of continuous remotely sensed data, and the radiometry of pixel values should be carefully considered before applying such measures. For instance, relying on float (decimal) precision data such as the NDVI (which ranges from −1 to 1), may lead to a high neighboring heterogeneity which could actually be the effect of data binning rather than the effect of an ecological underlying pattern. In general, an 8-bit image (and therefore composed of 256 integer values/classes) is preferable when applying spectral heterogeneity measures. In this paper, we used an 8-bit NDVI layer rescaled from Copernicus data. However, one might even rely on a multispectral system reduced to a single 8-bit layer through by means of the first component of a Principal Component Analysis or any multidimensionality reduction technique (Féret and Boissieu, 2020).

Most of metrics based on Information Theory can accommodate only one layer at a time when relying on indices using abundance information only (in the shown example, the Rényi entropy index and Shannon’s *H*). However, the `rasterdiv` package includes accounting for pixel values distances, such as the Rao’s *Q* index, which can integrate multiple layers of ecological information such as multiple bands of satellite images or physical and biotic data (e.g., vegetation cover, soil pH, topography).

In general, remotely sensed data are simplifications of more complex systems depending on the radiometric and spectral properties of one or more images. From an ecological point of view, the spectral space of an image might be associated with the Hutchinson’s hypervolume which orders geo-metrically the variables shaping species’ ecological niches (Hutchinson, 1959; Blonder, 2018). Hence, calculating heterogeneity in such a space could pro-vide important information about species niches variability or at least on the landscape variability shaping species distribution (Rocchini et al., 2018).

Future local and global changes are expected to impact ecosystem heterogeneity. Since remotely sensed data nowadays allows us to rely on relatively long and standardized time series, applying different measures of heterogeneity to multi-temporal stacks would enhance the power to estimate and potentially forecast ecosystem heterogeneity shifts in space and time. This will be an invaluable tool to allow targeted and efficient monitoring and planning practices. For instance, due to the unprecedented rate of climate change, the adaptation of species to climate change is a benchmark in ecology (Stein et al., 2014). The `rasterdiv` package might also be particularly useful when aiming at calculating climate-related heterogeneity, which is likely to shape ecosystem heterogeneity patterns that species have adapted to. This could be directly done running the functions on remotely sensed climate data (Metz et al., 2014; Senner et al., 2018; Zellweger et al., 2019), which are expected to drive several ecological functions at different spatial scales.

## 5 Conclusion

Measuring heterogeneity from space to understand ecological patterns and processes acting across the landscape and over different time periods is crucial to guide effective management practices, especially in the Anthropocene epoch, in which human intervention is leading to rapid environmental changes (Randin et al., 2020).

The proposed `rasterdiv` package is a powerful tool for monitoring spatial and temporal variation of ecosystems’ properties (Rocchini et al., 2018), given the intrinsic relationship (*sensu* Laliberté et al., 2019)between the spatial variation of ecosystems and that of the spectral signal from pixel values (Rocchini et al., 2019). No single measure can provide a full description of all the different aspects of ecosystem heterogeneity. That is why the `rasterdiv` package offers multiple approaches to disentangle the complexity of ecosystem heterogeneity in space and time through calculations deeply rooted in Information Theory and based on reproducible Free and Open Source algorithms.

## Acknowledgments

We are grateful to the handling Editor and two anonymous reviewers who helped us improving a previous version of this manuscript with their precious suggestions. DR and DK were partially supported by the H2020 project SHOWCASE. DR was also partially supported by the H2020 COST Action CA17134 “Optical synergies for spatiotemporal sensing of scalable ecophysiological traits (SENSECO)”. The research carried out at the Jet Propulsion Laboratory, California Institute of Technology, was under a contract with the National Aeronautics and Space Administration. Government sponsorship is acknowledged.

## Authors’ contribution to the manuscript

DR, MI, ET, MM, DDR, EM, CT, SV contributed to the development of the algorithms and the coding of the `rasterdiv` package.

DR, GB, MB, AB, GMF, RF, DK, SL, JL, MM, FM, AM, VM, BN, DP, CR, MR, FS, MJS, MS, LS, PS, AKS, SS, ET, PZ, MW contributed to the conceptual development of the theoretical background of the `rasterdiv` package.

All authors contributed to the writing of the manuscript.

## Data accessibility

Free data for applying the proposed functions are available directly into the `rasterdiv` package (https://CRAN.R-project.org/package=rasterdiv). The Copernicus Sentinel-2 image used in this paper can be freely downloaded from: https://scihub.copernicus.eu/, image tile reference: *S2A*_*M*_ *SIL*2*A*_2_0200905*T* 101031_*N*_ 0214_*R*_022_*T*_ 32*TPS*_2_0200905*T* 130252.

## Appendix 1 Mathematical proof of the equivalence between Rényi entropy and Shannon’s H given *α* → 1

### Theorem (De l’Hôpital).

*Let f, g* : (*a, b*) → ℝ *be two functions such that*

- lim_*x→a*_ *f* (*x*) = lim_*x→a*_ *g*(*x*) = 0
- *f and g are derivable in* (*a, b*) *with g*^*′*^ (*x*) ≠ 0 *for every x* ∈ (*a, b*)
- *the limit* 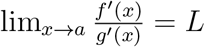 *with L* ∈ ℝ

*then*

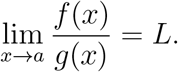

We now verify if the hypothesis hold in our case. Let *f* : (1, +∞) → ℝ and *g* : (1, +∞) → ℝ be functions such that 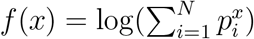 and *g*(*x*) =1 − *x* for every *x* ∈ (1, +∞). Since *p*_*i*_ is a fixed number in (0, 1) for every *i* ∈ {1, …,} with 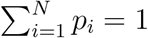, then

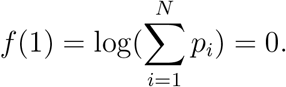

Moreover, since *f* is a composition of derivable functions in (1, + ∞), *f* is derivable in the definition domain. It is trivial that *g*(1) = 0, that *g* is derivable in the domain set and that *g*^*′*^(*x*) ≠ 0 for every *x*∈ (1, + ∞). The limit of the derivatives ratio is:

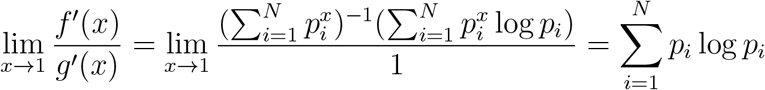

which is finite due to the hypothesis over *p*_*i*_. Since all the hypotheses of De l’Hôpital’s Theorem hold, the thesis follows, i.e.

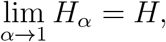

or Rényi ‘s entropy equals to Shannon’s *H* for *α* → 1.

